# Root Responses to Heterogeneous Nitrate Availability are Mediated by *trans*-Zeatin in Arabidopsis Shoots

**DOI:** 10.1101/242420

**Authors:** Arthur Poitout, Amandine Crabos, Ivan Petřík, Ondrej Novák, Gabriel Krouk, Benoît Lacombe, Sandrine Ruffel

## Abstract

Plants are subjected to variable nitrogen (N) availability including frequent spatial nitrate (NO_3_^-^) heterogeneity in soil. Thus, plants constantly adapt their genome expression and root physiology in order to optimize N acquisition from this heterogeneous source. These adaptations rely on a complex and long-distance root-shoot-root signaling network that is still largely unknown. Here, we used a combination of reverse genetics, transcriptomic analysis, NO_3_^-^ uptake experiments and hormone profiling under conditions of homogeneous or heterogeneous NO_3_^-^ availability to characterize the systemic signaling involved. We demonstrate the important role of the *trans*-zeatin form of cytokinin (CK) in shoots, in particular using a mutant altered for ABCG14-mediated *trans*-zeatin-translocation from the root to the shoot, in mediating: (*i*) rapid long distance N-demand signaling and *(ii)* long term functional adaptations to heterogeneous NO_3_^-^ supply, including changes in NO_3_^-^ transport capacity and root growth modifications. We also provide insights into the potential CK-dependent and independent shoot-to-root signals involved in root adaptation to heterogeneous N availability.

## INTRODUCTION

Continuous functional and morphological plasticity of organs is one of the most fascinating differences between plants and animals. Indeed, fixed in their environment, plants display a range of strategies allowing them to face fluctuating resource availability. Plant roots are particularly implicated since they are exposed to nutrient and water scarcity or over abundance. Signaling networks behind root adaptation are now central targets for the new green revolution aiming at optimizing belowground functioning to the benefit of aboveground development (Den Herder et al. 2010; Bishopp and Lynch 2015; Kong et al. 2014).

Consistent with the essential role of Nitrogen (N) for the biosynthesis of proteins, nucleic acids or essential pigments such as chlorophyll, roots are responsive to the availability of this element (Gruber et al. 2013; O’Brien et al. 2016; Kellermeier et al. 2014; Forde 2014). Responsiveness relies on the ability to sense N availability. This sensing is commonly divided into 2 main branches: local perception of N in the vicinity of the root (in particular the mineral N form nitrate, NO_3_^-^) and systemic perception of internal N/NO_3_^-^ availability at the whole organism level, relying on root-shoot-root signaling and integration of information in different parts of the plant (Walch-Liu et al. 2005; Gansel et al. 2001; Li et al. 2014; Alvarez et al. 2012). This dual sensing is integrated through an intricate signaling network permitting a mutual control of root N acquisition with plant growth to ensure N homeostasis (Krouk et al. 2011).

In *Arabidopsis*, perception and propagation of local NO_3_^-^ signaling has received a large attention. The molecular actors involved in this process include the NO_3_^-^ transceptor NPF6.3/NRT1.1/CHL1 (Ho et al. 2009; Krouk et al. 2010a), some kinases and phosphatase (CIPK8, CIPK23, ABI2, CPK10,30,32) (Hu et al. 2009; Ho et al. 2009; Liu et al. 2017; Léran et al. 2015) and several transcription factors (NLP6/7, TGA1/4, NRG2, SPL9) (Castaings et al. 2009; Marchive et al. 2013; Konishi and Yanagisawa 2013; Alvarez et al. 2014; Xu et al. 2016; Krouk et al. 2010b) targeting the expression of genes involved in NO_3_^-^ transport and assimilation, also known as the Primary Nitrate Response (PNR) (Medici and Krouk 2014). In addition, Ca^2+^ has been defined as a secondary messenger in this process (Riveras et al. 2015; Liu et al. 2017; Krouk 2017). Control of the N-response can be extended to additional transcription factors such as ANR1, ARF8 and NAC4 or CLE peptides that are involved in N-dependent root development (Zhang and Forde 1998; Vidal et al. 2013; Araya et al. 2014; Gifford et al. 2008) or bZIP1, LBD37/38/39 and BT2 transcription factors that control N-use (Rubin et al. 2009; Araus et al. 2016; Gutierrez et al. 2008). The TCP20 transcription factor, which is not involved in PNR *per se*, physically interacts with NLP6/7 to likely regulate the expression of NO_3_^-^ responsive genes and a cell cycle marker gene (Guan et al. 2014; Guan et al. 2017; Li et al. 2005). Interestingly, TCP20 is also a regulator of root foraging in heterogeneous N supply conditions (Guan et al. 2014) and thus could provide an anchorage point to understand how local and systemic N-regulation is integrated (Guan et al. 2017).

However, the functioning of a signaling cascade upstream of these local regulators of root physiology or morphology in response to the global N availability is poorly understood. This lack of knowledge is probably due to the necessity to use complex experimental approaches such as split-root system to address specifically the question of long distance signaling. Indeed, split-root experiments provide a relevant framework to untangle the different local and systemic signaling occurring in plants. The overall concept is to compare roots experiencing similar local hydro-mineral conditions but different distant (other part of the root) media (Li et al. 2014). By comparing roots in the same local condition, any differences between these roots can only be due to the impact of the long distance signal(s) (Figure 1A). This system helped to define the landscape of the N related systemic signaling response and defined at least 2 co-existing systemic signaling pathways (Ruffel et al. 2011; Li et al. 2014). The “N-demand” long distance signal conveys the information that the whole plant is experiencing a distal N-deprivation, while the “N-supply” signal conveys the information that some N has been found by the plant (Ruffel et al. 2011) (Figure 1A). Two components were identified in these signaling pathways: the role of C-terminally encoded peptides (CEP) (Tabata et al. 2014) and cytokinins (CK) biosynthesis (Ruffel et al. 2011; Ruffel et al. 2016).

The role of CEPs was recently demonstrated. Upon N-deprivation, CEPs are translocated to the shoots where they are recognized by the CEP Receptor 1 kinase (Tabata et al. 2014). Within the shoot vascular system, this recognition leads to the expression of small polypeptides that translocate toward the roots and participate, in combination with local NO_3_^-^, in controlling specifically the transcript accumulation of the main root high affinity NO_3_^-^ transporter *NRT2.1* (Ohkubo et al. 2017; Ruffel and Gojon 2017). However, in heterogeneous NO_3_^-^ supply conditions, roots display a wide range of adaptive responses including: the transcriptional regulation of hundred of genes, an enhanced lateral root development, and an enhanced N acquisition (Gansel et al. 2001; Remans et al. 2006; Ruffel et al. 2008; Ruffel et al. 2011; Mounier et al. 2014). The role of the CEP-derived long-distance signal remains to be demonstrated in these other aspects of the adaptive response to N heterogeneity.

It has been shown that CKs are synthesized in roots and translocated to the shoots in response to N provision, leading to the control of shoot growth (Takei et al. 2001; Takei et al. 2004; Sakakibara et al. 2006; Osugi et al. 2017). The crucial role of CK in root response to long distance signals was demonstrated (Ruffel et al. 2011). However, an important question remains concerning the role of CK in the shoots to actually trigger the root response in a systemic context. In other words, is active CK in the shoots a component of the long-distance signaling controlling at the same time molecular and physiological root response to NO_3_^-^ heterogeneity?

In this work, we demonstrate that shoot *trans*-zeatin (*tZ*) type CK is indeed an essential element of long-distance signaling that controls *(i)* transcriptional reprogramming of roots and shoots, *(ii)* root growth and *(iii)* NO_3_^-^ transport activity, in response to heterogeneous NO_3_^-^ conditions. By combining a genetic approach targeting modification of CK content and translocation with exhaustive measurements of CK forms and a shoot transcriptome analysis, we demonstrate that unbalanced root response to NO_3_^-^ provision relies on the integration in shoots of *tZ*.

## RESULTS

### Root to shoot cytokinin translocation controls transcriptional response to N-demand long distance signal

To investigate the role of CK root to shoot translocation in N-related systemic signaling, we characterized a mutant lacking the ABCG14 (ATP-Binding Cassette Transporter Subfamily G) transporter and thus impaired in delivering CK to the shoot (Ko et al. 2014; Zhang et al. 2014). To do so, wild-type (WT) and mutant plants were grown in a hydroponic split-root system. Our experimental framework consisted in 3 conditions of N provision, providing 4 different root samples (Figure 1A): *(i)* a homogeneous N-replete environment (C.KNO3: both compartments have 1 mM KNO_3_), *(ii)* a homogeneous N-deprived environment (C.KCl: both compartments have 1 mM KCl), and *(iii)* a heterogeneous split environment (Sp.KNO3/Sp.KCl: one compartment has 1 mM KNO_3_, and the other has 1 mM KCl). Any difference recorded between C.KNO3 and Sp.KNO3 samples is the signature of a N-demand long distance signal, while any difference recorded between Sp.KCl and C.KCl samples is the signature of a N-supply long-distance signal. This logic is applied throughout the whole manuscript (Figure 1A). This framework has been used to test the specific and quick response of sentinel genes to N-systemic signaling that were identified previously from a dynamical root transcriptomic analysis following plant transfer to homogeneous or heterogeneous conditions (Ruffel et al. 2011). Sentinel genes belong to important functions including the NO_3_^-^ assimilation pathway *(NiR, G6PD3, UPM1, FNR2)* and NO_3_^-^ transport systems *(NRT2.1* (Filleur et al. 2001) and its functional partner *NRT3.1/NAR2.1* (Yong et al. 2010)).

In WT plants, mRNA accumulation of the sentinel genes in Sp.KNO3 roots was higher compared to the homogeneous control condition C.KNO_3_, showing that Col-0 roots responded to NO_3_^-^ heterogeneous availability through a systemic N-demand signal (Figure 1B) (Ruffel et al. 2011). As expected, in these hydroponic split-root conditions, the isopentenyltransferase *ipt3,5,7* mutant, altered for CK biosynthesis (Miyawaki et al. 2006), displayed an alteration of the response to systemic N-demand (Figure 1B) (Ruffel et al. 2011). In the *abcg14* mutant, the expression level of sentinel genes was lower as compared to Col-0, but more importantly Sp.KNO3 roots did not display any significant stimulation of sentinel gene expression as compared to C.KNO3 control roots. This demonstrates that *abcg14* is also impaired in triggering the response to systemic N-demand signaling (Figure 1B). Therefore, CK root to shoot translocation could be essential for plant response to heterogeneous environment. It is noteworthy that for both mutants the expression level of sentinel genes is still responsive to the local NO_3_^-^ availability (C.KNO3 and Sp.KNO3 *versus* Sp.KCl, C.KCl; Figure 1B), showing that these mutants still preserve their ability to detect NO_3_^-^ *per se*.

**Figure 1.**
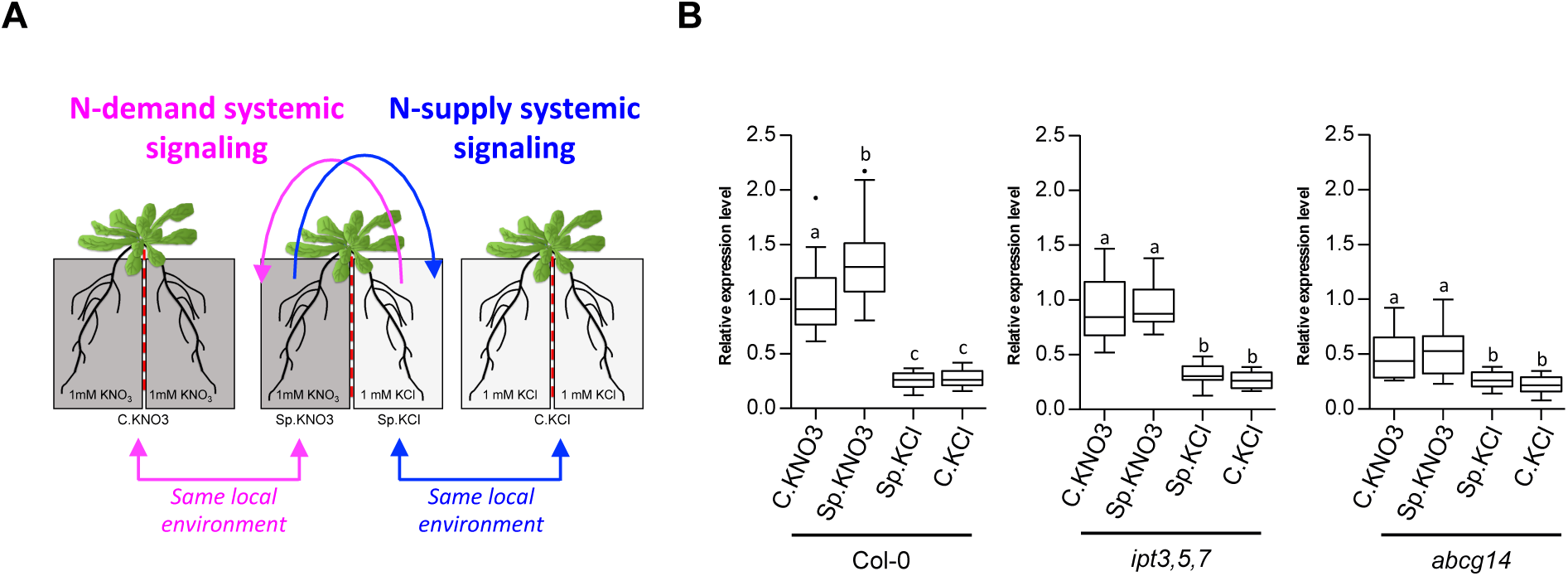
Perturbation of CK root to shoot translocation impairs the expression of N-demand sentinel genes. (**A**) WT and mutant plants were grown in hydroponic split-root conditions to decipher N-demand and N-supply signaling. (**B**) Relative expression level of sentinel genes was measured from roots harvested 6h30 after transfer in C.KNO3, Sp.KNO3/Sp.KCl and C.KCl conditions in Col-0, *ipt3,5,7* and *abcg14*. Boxplots display the expression values of the 6 N-demand sentinel genes *NRT2.1, NRT3.1, NiR, G6PD3, UPM1* and *FNR2*. Individual expression values have been normalized by the mean of the WT C.KNO3 expression across all experiments. Values are the means (+/− SE) of 3 independent experiments consisted each of 2 biological replicates corresponding to a pool of 3 plants. Different letters indicate significant difference according to one-way analysis of variance followed by a Tukey post-hoc test, p<0.05.

### Active *trans-zeatin* in shoots controls the root response to systemic N-demand

To determine how CK partitioning/homeostasis control the root specific reprogramming response to heterogeneous NO_3_^-^ supply, levels of the four basic isoprenoid CK types (*t*Z, isopentenyladenine iP, *cis*-zeatin *c*Z and dihydrozeatin DHZ) and their derivatives (ribotides, ribosides, *O*-glucosides and *N*-glucosides) were measured in roots and shoots, at the time point used to evaluate the response of sentinel genes (Figure 1).

#### CK partitioning is under the control of a combined effect of Nitrogen, IPTs and ABCG14

Firstly, as a control of our experimental system, we did a global analysis showing that the pattern of accumulation of the 4 CK types is indeed impacted by the *ipt3,5,7* and *abcg14* mutations as expected (Ko et al. 2014; Miyawaki et al. 2006) (Figure 2A). Moreover, we also provide insight into the response of CK accumulation to NO_3_^-^ provision in WT and these mutant genotypes (comparison C.KNO3 *versus* C.KCl; Figure 2A). As expected, the triple mutation in *IPT* genes led to a drastic decrease of *t*Z and iP-types in both shoots and roots (Figure 2A). In accordance with the predominant role of *IPT3* and *IPT5* in NO_3_^-^-dependent CK biosynthesis (Takei et al. 2004), NO_3_^-^ provision does not impact the low accumulation of *tZ* and *iP*-types still synthetized in the *ipt3,5,7* mutant (Figure 2A). Contrary to what has been previously observed, we did not see an increase of global *cZ*-type accumulation in the *ipt3,5,7* mutant (Miyawaki et al. 2006) but rather a significant decrease in the roots (Figure 2A). Interestingly, an increase of global *cZ*-type accumulation was rather observed in *abcg14* shoots in response to N-deprivation (C.KCl) (Figure 2A). A more detailed analysis of *c*Z-types revealed that in fact only *O*-glucosylated forms of *c*Z were increased in *ipt3,5,7* in all conditions (Supplemental Figure 1, blue arrows) whereas in *abcg14 O-*glucosides as well as transported and active *c*Z-forms were increased in shoots as soon as N provision was limited (Supplemental Figure 1, red arrows). Altogether, these results demonstrate that *c*Z-type homeostasis is indeed modified when MEP pathway-dependent CKs are perturbed. In addition, our experimental set-up provides an interesting framework to investigate the role of *c*Z-forms to maintain in shoots a minimal CK activity required to respond to abiotic stress (Schafer et al. 2015).

For the *abcg14* mutation, perturbations of CK partitioning and accumulation were consistent with previous data (Zhang et al. 2014; Ko et al. 2014). We indeed observed an increase in the accumulation of *t*Z-, iP- and DHZ-types in roots and a decrease in the accumulation of *t*Z-types in shoots, in accordance with the role of ABCG14 in root to shoot CK translocation (Figure 2A). Moreover, iP accumulation in the *abcg14* mutant is increased by N provision (Supplemental Figure 2, green arrows). Very interestingly, shoot iP content of the WT plants follows the level of root N provision and this aspect is very strongly affected by the *abcg14* mutation (Supplemental Figure 1, pink arrows). In more detail, this resulted in a lower accumulation of active iP-forms in C.KNO3 and a higher accumulation of all iP-forms in C.KCl in the *abcg14* mutant (Supplemental Figure 1, pink arrows). Therefore, taken together, these results demonstrate that the dynamic accumulation of *t*Z in shoots is under the control of the ABCG14 protein, and that this differential accumulation also controls N-responsive accumulation of iP-type CKs.

#### Shoot and not root *t*Z accumulation explains root sentinel gene expression

To explain the early response of sentinel genes to the systemic N-demand signaling and their perturbations in the mutant backgrounds (Figure 1B), we analyzed the accumulation of active and transported CK-forms (Base and Riboside) in roots and shoots in the split-root framework (Figure 1A). No obvious correlation was detected between gene expression (Figure 1B) and active-CK forms accumulation in roots (Figure 2B, left panel). Indeed, the *ipt3,5,7* and *abcg14* mutants display an opposite phenotype concerning *t*Z and iP accumulation whereas, in the same conditions, they both display the same gene expression profile (loss of N-demand signaling). We thus conclude that CK accumulation in roots cannot explain sentinel gene expression. However, shoot *t*Z accumulation can explain the root transcriptomic profile. Indeed, we observed that shoot *t*Z accumulation is decreased by both the *ipt3,5,7* and *abcg14* mutations (Figure 2B, right panel). In conclusion, the analysis of *ipt3,5,7* and *abcg14* mutants demonstrates that *t*Z accumulation in shoots is an explanatory factor of root gene expression in response to long-distance signaling.

Moreover, it is noteworthy that root *t*Z and iP differential accumulation between C.KNO3 Sp.KNO3 conditions are conserved in WT and the *abcg14* mutant (Figure 2B, left panel). This indicates a control of root CK accumulation by a systemic CK-independent N-demand signaling. This is consistent with our previous observations made on lateral root elongation harboring CK dependent and independent branches (Ruffel et al. 2016).

**Figure 2.**
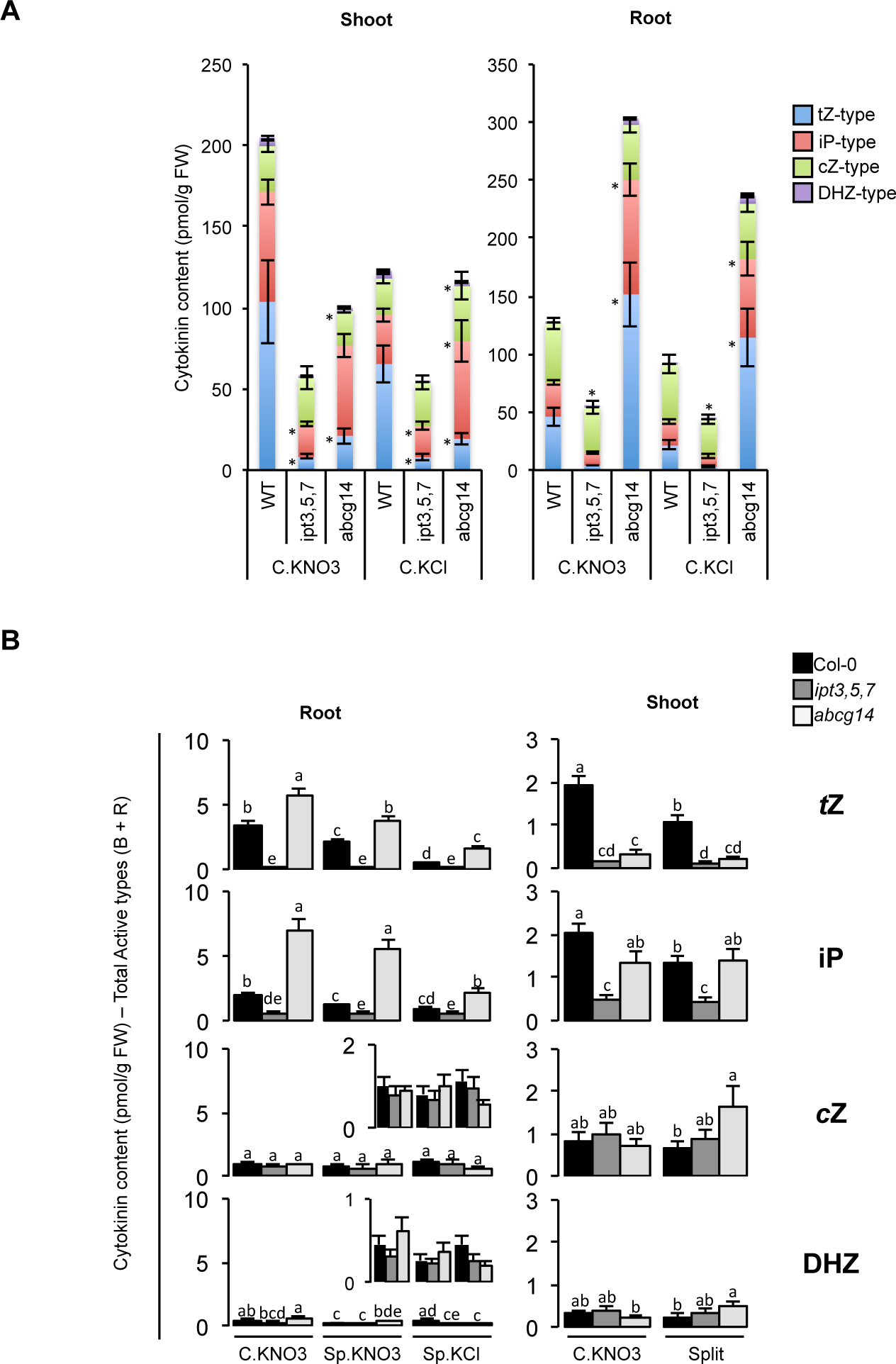
Root and shoot CK-types accumulation points on the role of *t*Z accumulation in shoots. (**A**) Stacked bar graphs display total CK accumulation and distribution of *tZ*, iP, *c*Z and DHZ type CK content in the shoots (left graph) and in the roots (right graph) from WT and mutant plants exposed to homogeneous C.KNO3 or C.KCl conditions. Asterisks indicate significant differences of accumulation between mutants and Col-0, according to a Student Test, p<0.05. If above the bar, the asterisk indicates that the accumulation of the 4 CK-types is different. (**B**) Barplots show active and transported (Base and Riboside) CK content (*t*Z, iP, *c*Z and DHZ) in the roots (left graphs) and in the shoots (right graphs) of WT and mutant plants in C.KN03 or split conditions.. Values are the means (+/− SE) of 5 to 6 biological replicates collected from 4 independent experiments. Letters indicate significant differences between treated genotypes. iP: *N^6^*-(Δ^2^-isopentyl)adenine; *t*Z: *tram-*Zeatin; *c*Z: *cis-*Zeatin; DHZ: Dihydro-zeatin.

### Shoot genetic reprogramming in response to heterogeneous NO_3_^-^ supply is perturbed in cytokinin biosynthesis and translocation mutants

As we previously demonstrated that *t*Z accumulation in shoots is crucial to explain root sentinel adaptation to long distance N-demand signaling, we decided to evaluate the impact of *ipt3,5,7* and *abcg14* mutations on the response of the shoot transcriptome in plants experiencing homogeneous or heterogeneous NO_3_^-^ supply (Figure 3A). To do so, *Arabidopsis* Gene1.1 ST Affymetrix array strips have been used (See Material and Methods for details on samples, arrays and data analysis). In the WT, 780 transcript clusters (probes equivalent) were significantly differentially accumulated between C.KNO3 and Split conditions. 422 were found to be up-regulated in Split conditions compared to the control C.KNO3 and 358 were regulated in the opposite direction (Supplemental Table 1). Only the 745 non-ambiguous AGIs were kept for the following analysis. Hierarchical clustering of their expression level in the 2 treatments (C.KNO_3_, Split) and the 3 genotypes revealed that the regulation in Col-0 by heterogeneous NO_3_^-^-provision was strongly affected in the 2 mutants (Figure 3A). Therefore, in addition to being impaired in *t*Z accumulation in response to N-supply, the 2 mutants undergo a perturbation of their capacity to reprogram gene expression in response to NO_3_^-^-supply, likely disrupting N-systemic signaling controlling root responses. Interestingly, some semantic terms were enriched within the annotation of these genes, indicating biological functions likely under the control of CK accumulation in shoots. Among the genes up-regulated in heterogeneous compared to homogeneous NO_3_^-^ condition, we found a significant enrichment of 2 interpro domains, ‘ipr000583’ and ‘ipr017932’, which both correspond to glutamine amidotransferase class-II domain found in 3 genes involved in glutamate synthesis (*i.e*., AT2G41220, AT3G24090, AT5G04140) (Figure 3B). Moreover, the overrepresentation of the ‘glutamine’ term was also found in 4 other genes annotated as glutamine amidotransferase class-I and glutamate-ammonia ligase *(i.e*., AT1G53280, AT3G53180, AT4G26900, AT4G30550) (Figure 3B). This result suggests that, even if NO_3_^-^ is the genuine signal to trigger N-demand systemic signaling (Ruffel et al. 2011), its heterogeneous supply triggers modification of an N assimilation pathway in shoots, in a CK-dependent manner. Similarly, we observed term enrichment among the genes down-regulated in heterogeneous compared to homogeneous NO_3_^-^ conditions, corresponding to the interpro domain ‘ipr006688’ found in 3 ADP-ribosylation factors *(i.e*., AT3G49860, AT5G14670, AT1G02440) and ‘duo1’ (or ‘germline-specific’) found in 2 genes annotated as C2H2 Zinc Finger proteins and HAPLESS 2 gene (*i.e*., AT4G35280, AT4G35700, AT4G11720) (Figure 3B).

By integrating the expression level of the whole genome data set (*i.e*., 3 genotypes and 2 treatments), we also identified 669 unique genes responding similarly to the NO_3_^-^ treatment in the shoots of the 3 genotypes (Supplemental Table 2). This corresponds to genes whose regulation is likely not related to CK-dependent long-distance signaling. Hierarchical clustering displayed a first level of classification based on the differential regulation in NO_3_^-^ heterogeneous versus homogeneous conditions (Supplemental Figure 3A). The semantic enrichment analysis, of those genes revealed only few meaningful terms, including, for example, the enriched terms ‘uba-like’ found in the annotation of 3 genes related to ubiquitination processes *(i.e*., AT2G17190, AT4G11740, AT5G50870) (Supplemental Figure 3B).

Altogether, this shoot transcriptomic analysis showed a massive and quick reprogramming of gene expression accompanying distinct *t*Z accumulation in response to NO_3_^-^-supply. Moreover, this allowed us to confirm the occurrence of CK-dependent and CK-independent branches of N systemic signaling (Ruffel et al. 2016) and to narrow down the biological pathways associated with the respective signals.

**Figure 3.**
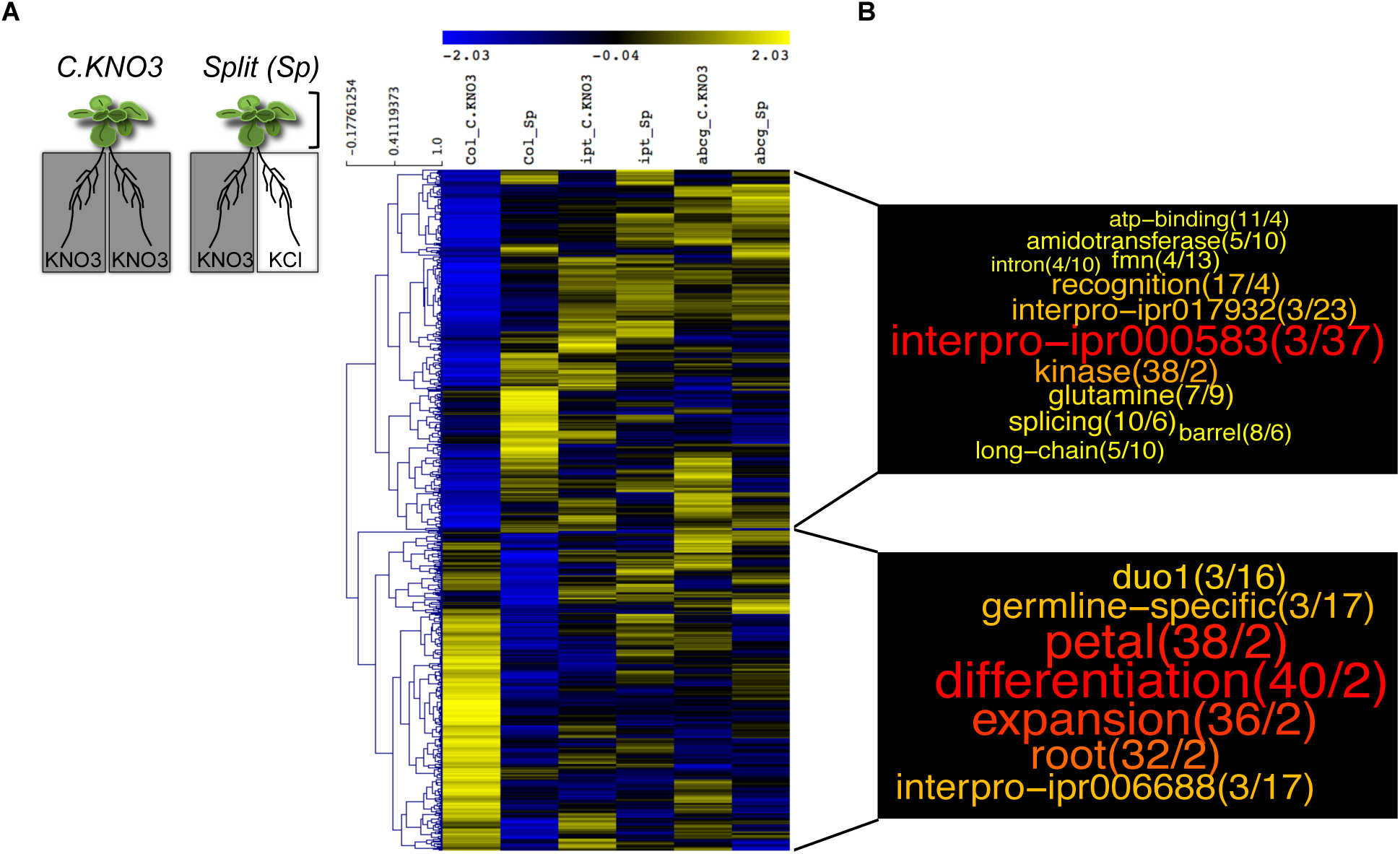
CK-dependent shoot transcriptome changes in response to root heterogeneous NO_3_^-^ supply. (**A**) Shoots of Col-0, *ipt3,5,7* and *abcg14* plants transferred for 24 hrs in NO_3_^-^ homogeneous (C.KNO3) or heterogeneous (Split) environment were harvested for transcriptomic analysis. Samples of 4 biological replicates from 4 independent experiments were used to perform the microarray analysis, using Arabidopsis Gene1.1 ST array Strip (Affymetrix GeneAtlas™). Hierarchical clustering of the 745 genes identified as differentially expressed in C.KNO3 versus Split conditions in the WT was performed with Multiple Experiment Viewer (MeV) software (http://mev.tm4.org/), using gene expression levels in the WT and the mutants. (**B**) Semantic enrichment in annotation of genes induced (at the top) or repressed (bottom) in Split condition compared to C.KNO3 in WT shoots, based on a GeneCloud analysis (https://m2sb.org). Beside each term, the first number corresponds to the number of genes containing the term and the second number gives the fold enrichment.

### *ipt3,5,7* and *abcg14* mutations affect integrated root traits in response to heterogeneous NO_3_^-^ supply

At a molecular level, sentinel genes responding specifically and quickly to a CK-dependent N-demand systemic signaling are largely involved in NO_3_^-^ transport and assimilation (*e.g*. NO_3_^-^ transporter NRT2.1 or Nitrite Reductase) (Figure 1B). Therefore, we asked to what extent genetic perturbation of CK biosynthesis and root to shoot translocation could affect the associated long-term adaptation of root physiology to different N supply: root NO_3_^-^ influx capacity and biomass (*e.g*. root dry weight). We also decided to highlight the relationship between these two components of N acquisition. To do so we combined root traits on single graphs allowing us to take into account the variability of plant growth between independent biological replicates (Figure 4A).

In WT, both components (root dry mass and NO_3_^-^ transport) responded significantly to N-systemic signaling (N-demand and N-supply) since the mean values are significantly higher in Sp.KNO3 (5.8 mg and 92 μmol N.h^−1^.g^−1^ root DW) compared to C.KNO3 (3.6 mg and 56 μmol N.h^−1^.g^−1^ root DW) and higher in C.KCl (4.3 mg and 74 μmol N.h^−1^.g^−1^ root DW) compared to Sp.KCl (2.9 mg and 28 μmol N.h^−1^.g^−1^ root DW) (Figure 4A, graphs on top). More importantly, this analysis revealed the existence of a relationship between the two adaptive processes in NO_3_^-^ supply roots (*i.e*. transport and biomass). Indeed, we observed that WT plants having a strong developmental response display a lower NO_3_^-^ transport adaptation and *vice-versa* (Figure 4A, top left).

In the mutants, root biomass and NO_3_^-^ influx responses to N-demand systemic signaling were deeply perturbed (Figure 4A, left panel), suggesting that CK and more precisely root to shoot translocation is a limiting factor for root response to a heterogeneous environment. The stimulation of root biomass production in Sp.KNO3 compared to C.KNO3 was abolished (Figure 4A, left panel). According to the strategy of WT plants described above, one would expect that the smallest root systems would display a greater increase in NO_3_^-^ influx as an alternative strategy to compensate for distal N-deprivation. Thus, the mutants might compensate for their root biomass phenotype by increasing their NO_3_^-^ transport activity. However, this is not what we observed. Indeed, if we consider for instance only roots with a dry biomass below the arbitrary value of 2 mg (expected to be the one to adapt the most their NO_3_^-^ transport), WT Sp.KNO3 roots displayed a NO_3_^-^ influx significantly 2 times greater than their respective controls whereas mutant Sp.KNO3 and C.KNO3 roots display the same low level of NO_3_^-^ influx (Figure 4B). Therefore, CK appears to be central to set up the two intertwined adaptive responses to heterogeneous NO_3_^-^ supply (*i.e*. NO_3_^-^ transport and root development).

Interestingly, the two mutants did not behave similarly in response to the N-supply systemic signaling (Figure 4A, right panel). Whereas CK biosynthesis is required to stimulate root proliferation and NO_3_^-^ influx when N is completely absent from the medium (Figure 4A, graph on middle right), the *abcg14* mutant still maintained a NO_3_^-^ influx capacity similar to Col-0 (*i.e*. 71 and 74 μmol N.h^−1^.g^−1^ root, respectively) albeit the *abcg14* mutant displayed a higher variability than WT (Figure 4A, graphs on top and bottom right).

Altogether, our results show that CKs have a broad role in regulating integrated adaptive traits in response to long-distance signaling pathways. In the same way, the modulation of an even more integrative trait controlled by N provision, that is the shoot/root ratio, is lost upon CK perturbation (Supplemental Figure 4).

We conclude that CKs are deeply involved in the control of long-term plant adaptation to NO_3_^-^ heterogeneity and that CK-translocation cannot explain the entirety of these responses since it’s likely not the root to shoot *t*Z translocation that is involved in regulating root NO_3_^-^ uptake capacity by N-supply systemic signaling.

**Figure 4.**
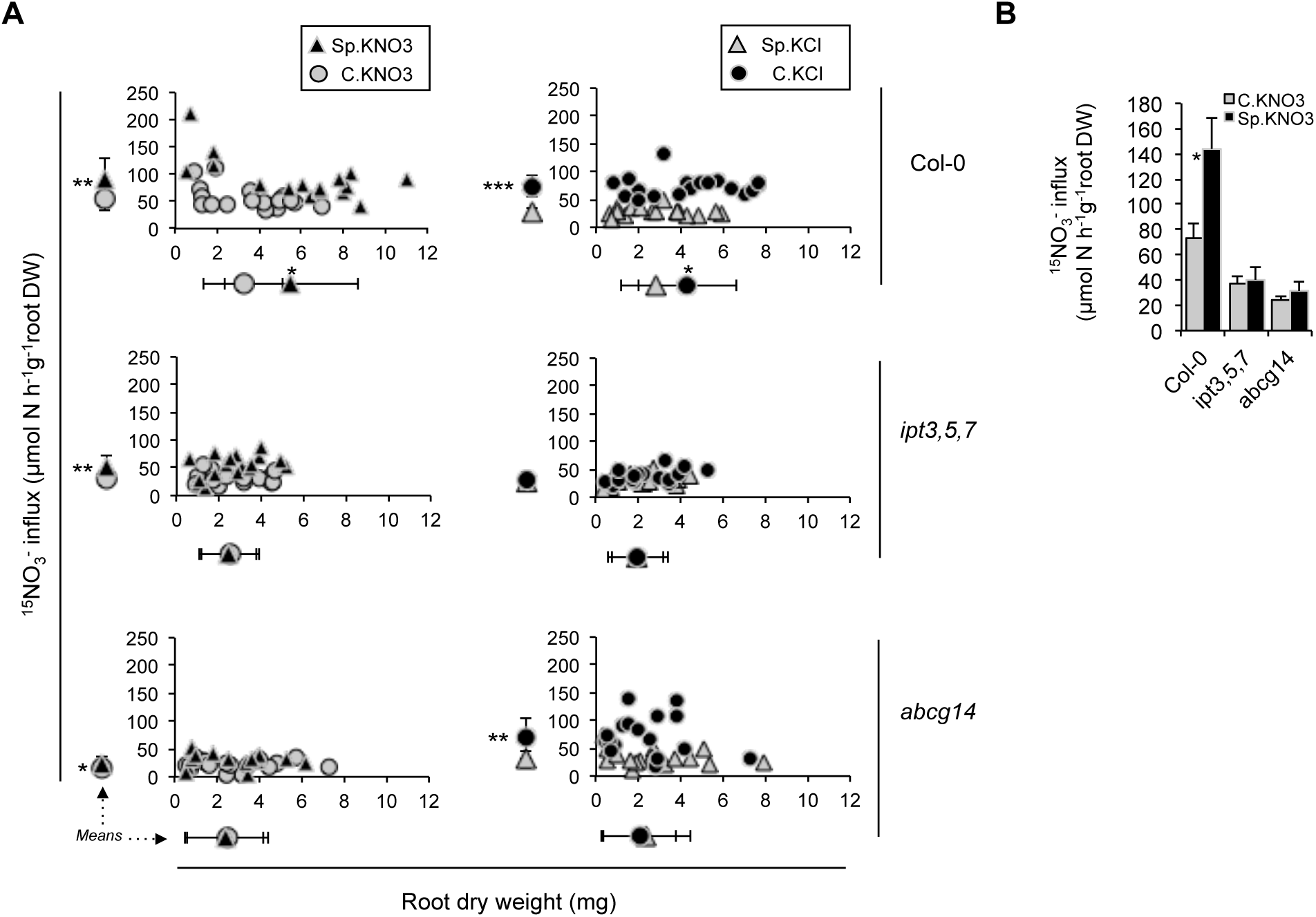
Responses of integrated traits to N-systemic signaling are affected in CK mutants. (**A**) Individual root ^15^NO_3_^-^ influx is plotted against the root biomass. Root biomass and root ^15^NO_3_^-^ influx means are presented out of the graphs, below and on the left of each plot, respectively. On the left panel, the effects of N-demand systemic signaling in Col-0, *ipt3,5,7* and *abcg14* are shown by comparing C.KNO3 and Sp.KNO3 roots. On the right panel, effects of N-supply systemic signaling in Col-0, *ipt3,5,7* and *abcg14* are shown by comparing C.KCl and Sp.KCl roots. Graphs on top, middle and bottom correspond to Col-0, *ipt3,5,7* and *abcg14*, respectively. Data were obtained from 3 independent experiments, each including 6 biological replicates. Asterisks indicate significant differences between means, according to t-test with p<0.05*, p<0.01**, p<0.001***. (**B**) To generate the bar graph, we used root NO_3_^-^ influx measurements from roots displaying dry weight below the arbitrary value of 2 mg, corresponding to 1/3 of the measurements presented in (A).

### Local nitrate response is not impaired in cytokinin mutants

Previously, we have shown that root response to heterogeneous NO_3_^-^ supply depends on a long-distance signaling network triggered by NO_3_^-^ *per se*. Indeed, the regulation of the specific and early sentinel genes in heterogeneous conditions is similar between WT plants and NR-null mutant, leading to the conclusion that the perception of NO_3_^-^ is a prerequisite to trigger the response to systemic signaling (Ruffel et al. 2011). Thus, in order to rule out that the *ipt3,5,7* and *abcg14* mutants are impaired for their molecular and physiological response to heterogeneous NO_3_^-^ conditions only because they can not perceive local NO_3_^-^ provision, we evaluated their respective responsiveness to homogeneous NO_3_^-^ supply. We first tested the PNR (Medici and Krouk 2014) in the two mutants, by transferring plants from N free solution to 1 mM KNO3 or 1 mM KCl for 30 min. In the two mutants, the activation of NO_3_^-^-responsive marker genes was similar to that observed in WT plants (Figure 5A). Although not significant, a reduction of NO_3_^-^ induction was observed for some nitrate-responsive marker genes in the *abcg14* mutant (e.g., *Nitrite Reductase-NiR)*. However, multiple biological replicates did not confirm that this mutant is impaired in NO_3_^-^ perception (Supplemental Figure 5). The capacity of *ipt3,5,7* and *abcg14* mutants to react to NO_3_^-^ addition was also supported by a similar root high affinity NO_3_^-^ influx level between the 3 genotypes, at the same time point selected to evaluate PNR (Figure 5B).

Similarly, the absence of root biomass changes in response to systemic signaling in the mutants (Figure 4A) prompted us to verify that they are not only restrained in their capacity to grow, independently of N supply conditions. When the three genotypes were grown on N-free, 0.1mM KNO3 or 1mM KNO3 containing media, the primary root length and total lateral root length increased with N concentration for the 3 genotypes (Figure 5C), with a longer primary root in the *ipt3,5,7* mutant as previously shown (Miyawaki et al. 2006). Thus, we confirmed that these mutants have the capacity to grow with increasing N concentration.

Altogether, we conclude that CK biosynthesis and root-to-shoot translocation do not impair NO_3_^-^ perception as well as the potential to use N or to grow continuously in accordance with N supply but is rather important to finely tune functional root response when N is partially or completely absent.

**Figure 5.**
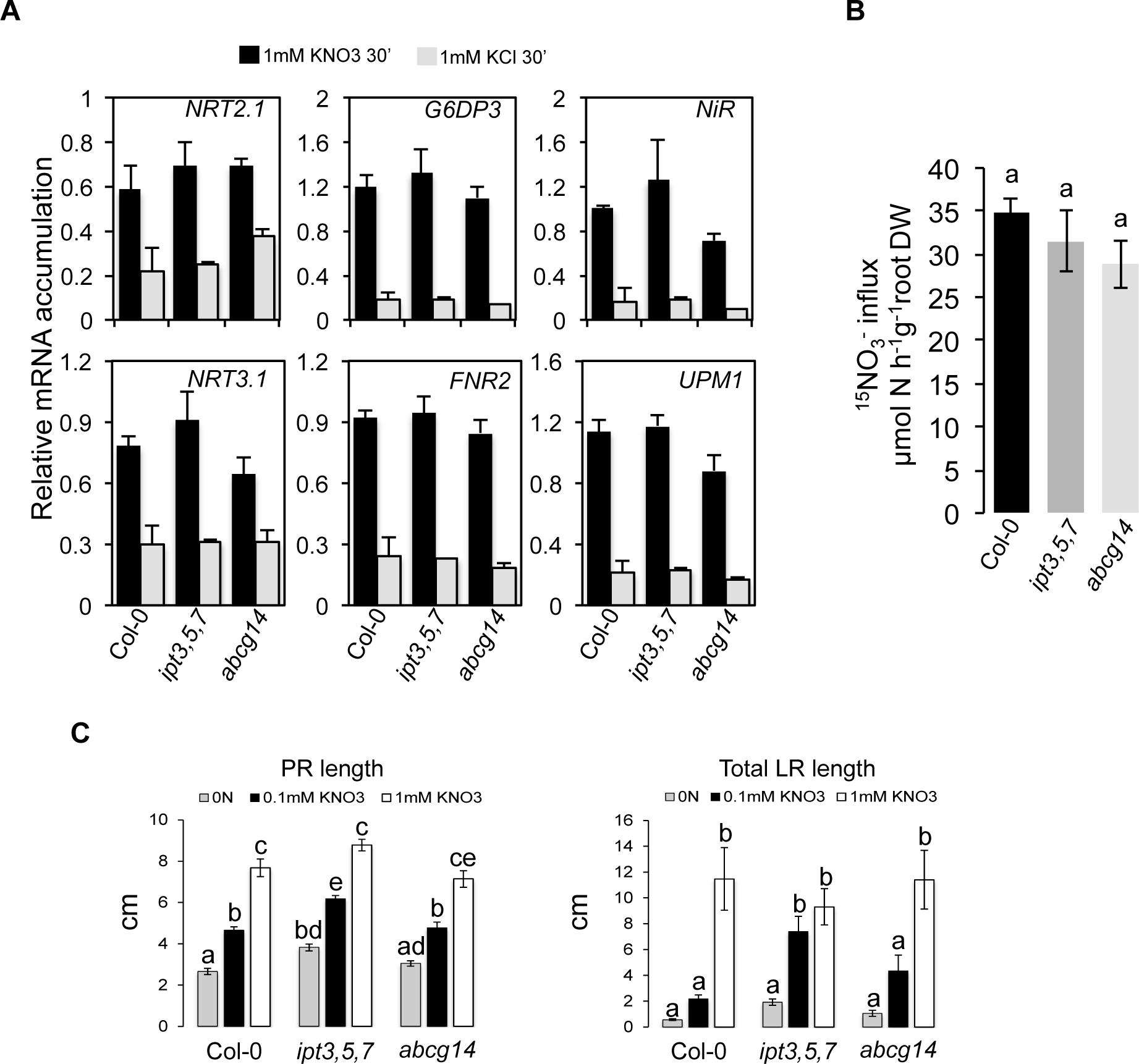
Primary NO_3_^-^ response and NO_3_^-^ dependent root development are not impaired in CK mutants. (**A**) Relative expression level of 6 marker genes of the primary NO_3_^-^ response *NRT2.1, NRT3.1 NIR, G6PD3, UPM1* and *FNR2* in roots from Col-0, *ipt3,5,7*, and *abcg14* plants transferred 30 min in media containing 1mM KNO_3_ (dark bars) or 1mM KC1 (grey bars) as the control. Data are means (+/− SE) obtained from 2 independent experiments including in each 2 pools of 5–6 plants. (**B**) Root ^15^NO_3_^-^ influx in 0.2 mM K^15^NO_3_, in Col-0 (dark bar), *ipt3,5,7* (dark grey bar) and *abcg14* (light grey bar) plants prior exposed to KNO_3_ 1 mM for 30 min. Data are means (+/− SE) obtained from 24 individual plants from 2 independent experiments. (C) Primary and lateral root length were measured in WT and mutant plants grown for 17 days in N-free, KNO_3_ 0.1 mM or KNO_3_ 1mM containing medium. Data are means (+/−SE) determined from 6 to 12 plants. Letters indicate significant difference according to a one-way analysis of variance followed by tukey post-hoc test; p<0.05.

## DISCUSSION

Functional root responses to N availability are the results of a complex signaling network integrating a localized sensing of root NO_3_^-^ availability with long-distance signaling aimed at coordinating the needs of the different parts of the plant. Here, we show that the integration of *t*Z content in shoots is an essential component of the long-distance signaling network controlling root responses. Our results, supported by previous works focusing on the functional characterization of genes involved in CK biosynthesis (Sakakibara et al. 2006; Kiba et al. 2013; Osugi et al. 2017), suggest that NO_3_^-^ triggers *t*Z biosynthesis mainly in roots. The *t*Z-types would be then transported to the shoots *via* ABCG14 where they modify gene expression and possibly the associated metabolism. Therefore, our model would propose that NO_3_^-^ provision lead to *t*Z accumulation in roots that are subsequently transported to the shoots. The accumulation of *t*Z, differing between homogeneous or heterogeneous conditions, would be integrated at the shoot level leading to a differential control of root response according to NO_3_^-^ provision applied to the roots. In this scenario, *t*Z translocation could even constitute a part of the systemic signal it-self triggering a shoot to root signal that still needs to be identified.

In shoots, glutamine biosynthesis is a semantic term significantly influenced by *t*Z accumulation (Figure 3). Interestingly, amino acids have been hypothesized to be reporters of the N status of the plant (Cooper and Clarkson 1989; Muller and Touraine 1992). We now hypothesize that CK-dependent NO_3_^-^ related signals could modify shoot glutamine or glutamate metabolism and that this might be part of a branch of the shoot-to-root signal as previously hypothesized by others (Imsande and Touraine 1994; Gent and Forde 2017; Girin et al. 2010). Of course, further studies will be necessary to validate this hypothesis, but our work provides new experimental and genome-wide clues concerning the potential role of amino acids as part of the shoot-to-root signaling.

We confirmed that CK accumulation is deeply affected in the *ipt3,5,7* and *abcg14* mutants (Figure 2) (Miyawaki et al. 2004; Ko et al. 2014; Zhang et al. 2014). Despite the strong repressive effect of local CK status on root development (Werner et al. 2010; Laplaze et al. 2007), we have found that these mutants, in particular *abcg14*, still maintain a certain capacity to adapt their development in particular to homogenous N provision (Figure 5C). This could be explained by recent results obtained on CK partitioning between the apoplasm and the cytosol. Indeed, an important aspect of CK signaling is that CKs are perceived in the apoplasm and that the PUP14 transporter is crucial to import bioactive CKs into the cytosol and suppress the CK response (Zürcher et al. 2016). In this perspective, we think that CK accumulation is important but active CK transport at the cellular level in roots can, to some extent, explain the responsiveness of *ipt3,5,7* and *abcg14* mutants.

Finally, this work refines the model of integration of the different N-related long-distance signaling pathways displaying some differences with the CEP-related long distance model. Indeed, CEPs were shown to be synthetized in N deprived roots and transported to the shoots where they are recognized by their related receptor kinase (CEPR1) (Tabata et al. 2014). This recognition activates the biosynthesis of CEPD1 polypeptides that are transported to the root where they activate *NRT2.1* transcript accumulation (Ohkubo et al. 2017). Thus the CEP model is drastically triggered by N deprivation. In our model, we believe that the long distance signal is generated by NO_3_^-^ itself. Indeed, we observed that *t*Z accumulation follows the NO_3_^-^ provision level (Figure 2B). We thus hypothesize that the 2 models are likely relying on different signaling modules. In nature, a plant shoot, experiencing root heterogeneous N conditions, likely receives a combination of different long-distance signals coming from the different parts of the plant (including CEP from N deprived roots and CK from NO_3_^-^ supply roots). Future investigations will aim to resolve how plants integrate in shoots these different signaling pathways to reach a coherent root adaptive response.

## MATERIALS AND METHODS

### Plant materials

*Arabidopsis thaliana* Col-0 background was used as WT plant. The *abcg14* (SK_15918) mutant line was kindly provided by Donghwi Ko (The Sainsbury Laboratory, Cambridge, UK). The *ipt3,5,7* triple mutant line was previously kindly provided by Sabrina Sabatini (University “La Sapienza”, Rome).

### Plant growth conditions

All plants were grown in short day light period (8h light 23°C/16h dark 21°C) at 260 μmol.m^−2^.s^−1^ intensity. Split-root *in vitro* culture was done as previously described (Ruffel et al. 2011). Briefly, plants were grown on solid (1% agar type A) N-free modified basal MS medium complemented with 0.5 mM NH_4_-succinate and 0.1 mM KNO_3_ as N sources. At day 10, the primary root was cut off below the second lateral root, to obtain 2 news ‘primary roots’. At day 14, plants were transferred in 1 mM NH_4_-succinate splitted-medium in order to separate the root system in two isolated parts. At day 18, plants were transferred in new split plates containing: basal MS medium supplemented with 1 mM KNO_3_ on one side (Sp.KNO_3_) and 1 mM KCl on the other side (Sp.KCl) or 1 mM KNO_3_ on both side (C.KNO_3_) or 1 mM KCl on both side (C.KCl). Mini hydroponic *in vitro* culture (Phytatrays) was based on the same media described above and a similar timing but it did not include any root pruning. For split-root in hydroponic system, seeds were sown on upside down eppendorf taps with 1 mm whole filled by H_2_O-agar 0.5% solution and grown during 7 days on tap water. Then, seedlings were grown on nutritive solution containing KH_2_PO_4_ 1 mM; MgSO_4_,7H_2_O 1 mM; K_2_SO_4_ 0.25 mM; CaCl_2_,2H_2_O 2.5 mM; Na-Fe-EDTA 0.1 mM; KCl 50 μM; H_3_BO_3_ 7.5 μM; MnSO_4_,H_2_O 1.25 μM; ZnSO_4_,7H_2_O 0.25 μM; CuSO_4_, 5H_2_O 0.25 μM; (NH_4_)6 Mo_7_ O_24_, 4H_2_O 0.025 μM; supplied with 1 mM NH_4_Cl and 0.1 mM KNO_3_ as N sources, pH 5.8. Nutritive solution was renewed every 4 days. 17 days after sowing, primary root was cut off below the second lateral root, to obtain 2 root systems. After 4 additional days, plants were transferred in split root system with 1 mM NH4Cl as the sole N source for 4 more days to let the roots grow in split conditions. 24 hrs before treatment, nutrient solution is renewed by a N-free nutritive solution. Treatments are applied by adding concentrated KNO3 or KCl solution in each compartment up to a final concentration of 1 mM. For PNR analysis, plants were grown exactly as we did for split-root experiments, except that the primary root was not cut and thus the root system was not splitted in 2 parts at the time of the treatment. For root development traits analysis, plants were sown and grown for 17 days *in vitro*, in square plates containing modified N-free basal MS supplied with 0, 0.1 or 1 mM KNO_3_, 0.3 mM sucrose, 0.5 g/L MES and 1% agar type A. Time collection or analysis of plants tissues was done as indicated in the results.

### Gene expression analysis

Total RNA was extracted from frozen and grounded root or shoot tissues using TRIzol^TM^ reagent (15596026, ThermoFisher Scientific), following provider’s instructions. RNA integrity and concentration were determined using a 2100 Bioanalyzer Instrument (Agilent) and Agilent RNA 6000 Nano kit (5067–1511, Agilent). DNA contamination was removed by digestion with DNase I (AMPD1, SIGMA). For real-time qPCR analysis, reverse transcription of mRNAs was done using ThermoScript™ RT-PCR (11146016, ThermoFisher Scientific) according to the manufacturer’s protocol. Gene expression was determined using a LightCycler^®^ 480 Instrument (Roche) and SYBR^®^ Premix Ex Taq^TM^ (RR420L, TaKaRa). Expression levels of tested genes were normalized using the expression level of *Actin2/8* and *Clathrin* genes. All specific primers used in this study are listed in Supplemental Table 3. Genome-wide expression analysis in shoots was based on 4 biological replicates obtained from 4 independent experiments including the 3 treatments (*i.e*., C.KNO_3_, Split, C.KCl) and the 3 genotypes (*i.e*., Col-0, *ipt3,5,7, abcg14*). Gene expression measurements were performed using *Arabidopsis* Affymetrix^®^ Gene1.1 ST array strips designed to measure whole transcript accumulation of 28.501 genes (or transcripts clusters), based on 600.941 probes defined on TAIR10 genome annotation. Biotin labeled and fragmented cRNAs were obtained using GeneChip^®^ WT PLUS Reagent kit (902280, ThermoFisher Scientific) following manufacturer’s instructions. Hybridization on array strips was performed for 16 hours at 48°C. Arrays are washed, stained and scanned using GeneAtlas HWS Kit (901667, ThermoFisher Scientific) on the GeneAtlas^®^ Fluidics and Imaging Station.

### Statistical analysis and Bioinformatics

Microarrays raw data were processed with GCRMA available on the Expression Console Software developed by Affymetrix. Data analysis was performed in [R]. Genes differentially expressed specifically in WT were identified by a t-test analysis (p-value<0.05). Genes responding to the treatment in 3 genotypes have been identified using a two-way ANOVA that was modeled as follows: Y= μ + α_genotype_ + β_treatment_ + (αβ)_genotype_* _treatment_ + ε, where Y is the normalized expression signal of a gene, μ is the global mean, the *α* and β-coefficients correspond to the effects of NO_3_^-^ availability (homogeneous or heterogeneous), of the genotype and of the interaction between both factors, and ε represents unexplained variance. All the genes for which at least the β_treatment_ is significant (p-value<0.05) to explain variation of expression have been selected. Hierarchical clustering of gene expression was performed using MultiExperiment Viewer v4.8 (MeV) software (Saeed et al. 2003). Functional analysis of gene lists was performed using the GeneCloud platform and semantic enrichment was displayed using word clouds (https://m2sb.org) (Krouk et al. 2015).

### Determination of cytokinin content

CK purification was performed according to the described method (Svačinová et al. 2012) with modifications (Smehilova et al. 2016). Briefly, CKs were extracted from 30 mg of frozen powder in modified Bieleski buffer (methanol/water/formic acid, 15/4/1, v/v/v) together with a cocktail of stable isotope-labeled internal standards (0.25 pmol of CK bases, ribosides and *N*-glucosides, 0.5 pmol of CK *O-*glucosides and nucleotides per sample added), and purified using two solid phase extraction columns. CK content was determined by UHPLC–MS/MS (Ultra-High Performance Liquid Chromatography coupled to a triple quadrupole mass spectrometer equipped with an electrospray interface). The quantification was performed by Masslynx software (v4.1; Waters) using a standard isotope dilution method. The ratio of endogenous CK to the appropriate labeled standard was determined and further used to quantify the level of endogenous compounds in the original extract according to the known quantity of the added internal standard.

### Determination of root biomass and nitrate influx capacity

Root ^15^NO_3_^-^ influx was assayed as described previously (Munos et al. 2004). Root systems were rinsed with 0.1 mM CaSO_4_ solution for 1 min, transferred to nutrient solution containing 0.2 mM ^15^NO_3_^-^ (99% atom exess ^15^N) at pH 5.8, for 5 min and washed with 0.1 mM CaSO_4_ solution for 1 min. Roots and the shoots were harvested separately and dried in a oven at 70°C for 48 hrs. Dry weight was determined and the total nitrogen and atom % ^15^N were determined by continuous-flow isotope ratio mass spectrometer, using a Euro-EA Euro Vector elemental analyzer coupled with an IsoPrime mass spectrometer (GV Instruments).

### Measurements of root development traits

Scans of the square plates containing the plants were performed at 600 dpi in TIFF format using a HP scanner. Length of primary and lateral roots as well as the number of lateral roots were measured using ImageJ software (Rasband 1997–2016).

## AUTHOR CONTRIBUTIONS

A.P., A.C., I.P. and O.N. performed research;. A.P. and S.R. analyzed data. G.K., B.L. and S.R. designed research and wrote the manuscript.

## ACKNOWLEDGEMENTS

We thank Hugues Baudot and his team for taking care of the plant culture system, Pascal Tillard for ^15^N measurements and Hana Martínková for her help with phytohormone analyses.

## Supplemental Information

1. Supplemental Figures 1 to 5

2. Supplemental Tables 1 to 3 (uploaded in separate files).

**Supplemental Figure 1.**
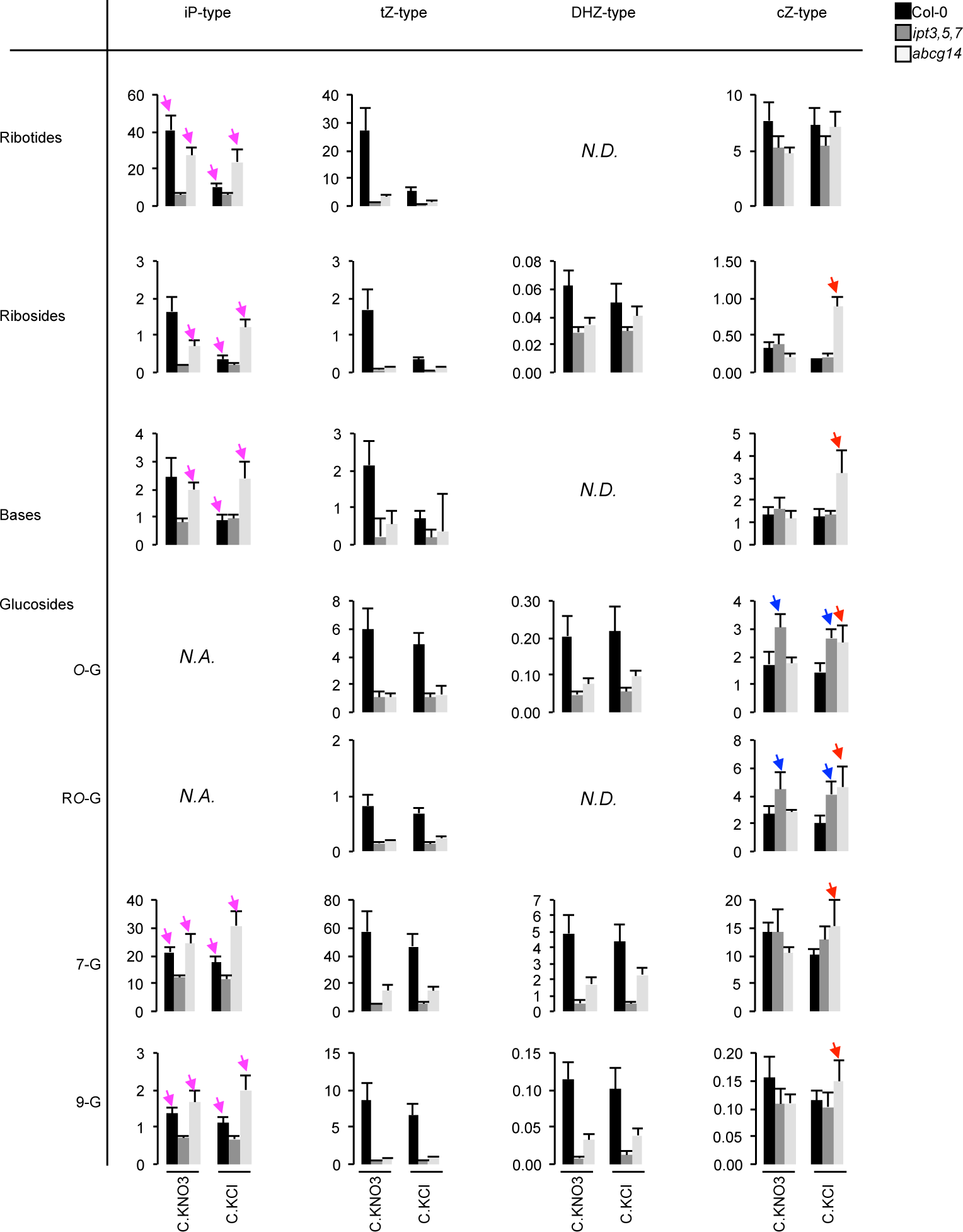
CK contents in shoots of WT, *ipt3,5,7*, and *abcg14* plants exposed to split-root conditions. Barplots show total shoot accumulation of *tZ*, iP, *cZ* and DHZ-type CK respectively from left to right and ribotides, ribosides, bases and glucosides forms respectively from top to bottom. Black bars: WT; dark grey bars: *ipt3,5,7*; light grey bars: *abcg14*. Values are the means (+/− SE) of 5 to 6 biological replicates collected from 4 independent experiments. N.D.: Not Detected. N.A.: Not Applicable. iP: *N*^6^-(Δ^2^-isopentyl)adenine; *t*Z: *trans*-Zeatin; *c*Z: *cis*-Zeatin; DHZ: Dihydrozeatin. Pink arrows indicate that N-dependent iP accumulation is affected in *abcg14* compared to Col-0. Blue arrows indicate an increase of *O*-glucosylated *c*Z in *ipt3,5,7*. Red arrows indicate increase of *c*Z in *abcg14* in N-deprived conditions.

**Supplemental Figure 2.**
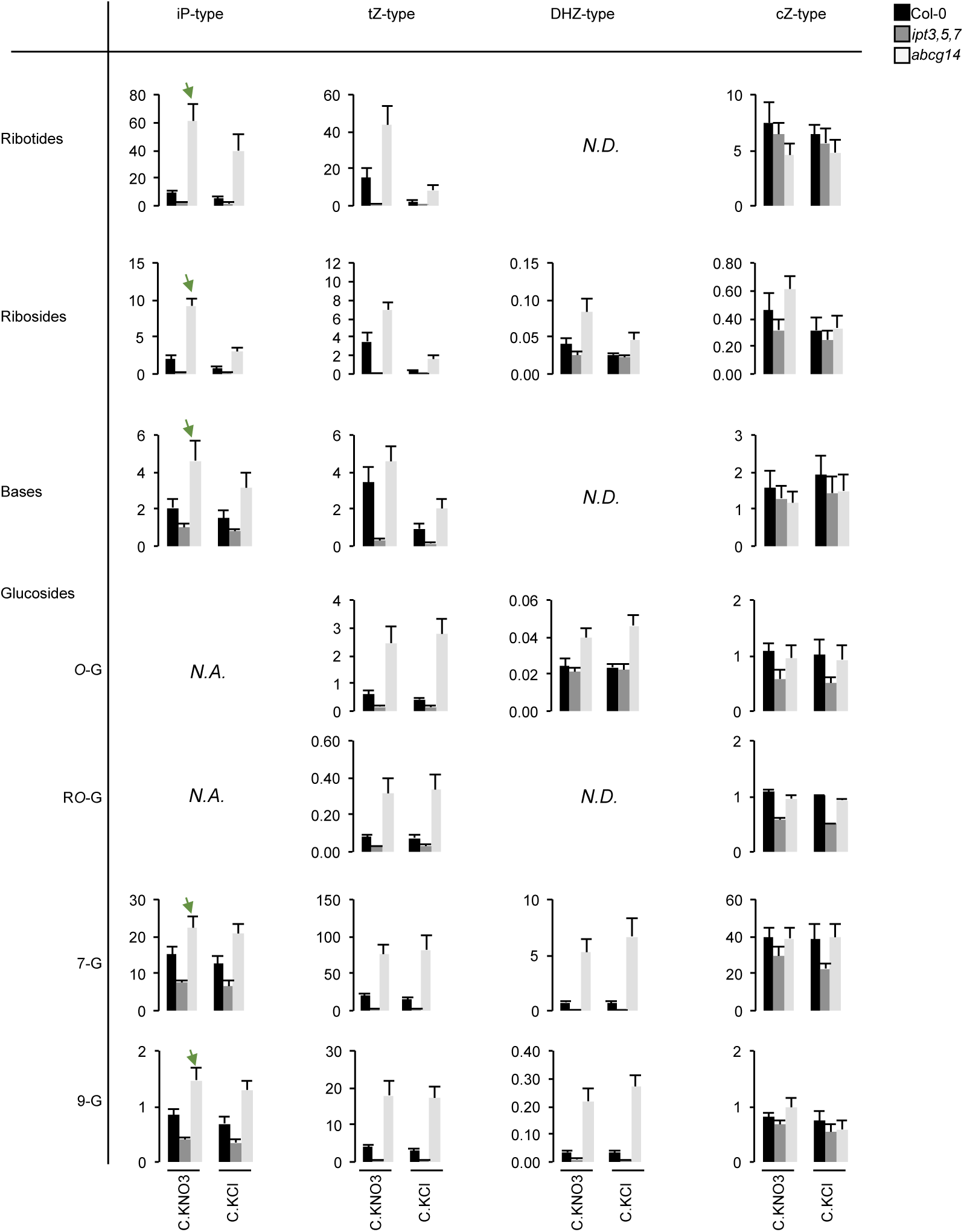
CK contents in roots of WT, *ipt3,5,7*, and *abcg14* plants exposed to C.KNO3 or C.KCl conditions. Barplots show total root accumulation of *tZ*, iP, *cZ* and DHZ-type CK respectively from left to right and ribotides, ribosides, bases and glucosides forms respectively from top to bottom. Black bars: WT; dark grey bars: *ipt3,5,7*; light grey bars: *abcg14*. Values are the means (+/− SE) of 5 to 6 biological replicates collected from 4 independent experiments. N.D.: Not Detected. N.A.: Not Applicable. iP: *N*^6^-(Δ^2^-isopentyl)adenine; *tZ: trans-*Zeatin; *c*Z: *cis*-Zeatin; DHZ: Dihydrozeatin. Green arrows indicate N-dependent iP accumulation in *abcg14* mutant.

**Supplemental Figure 3.**
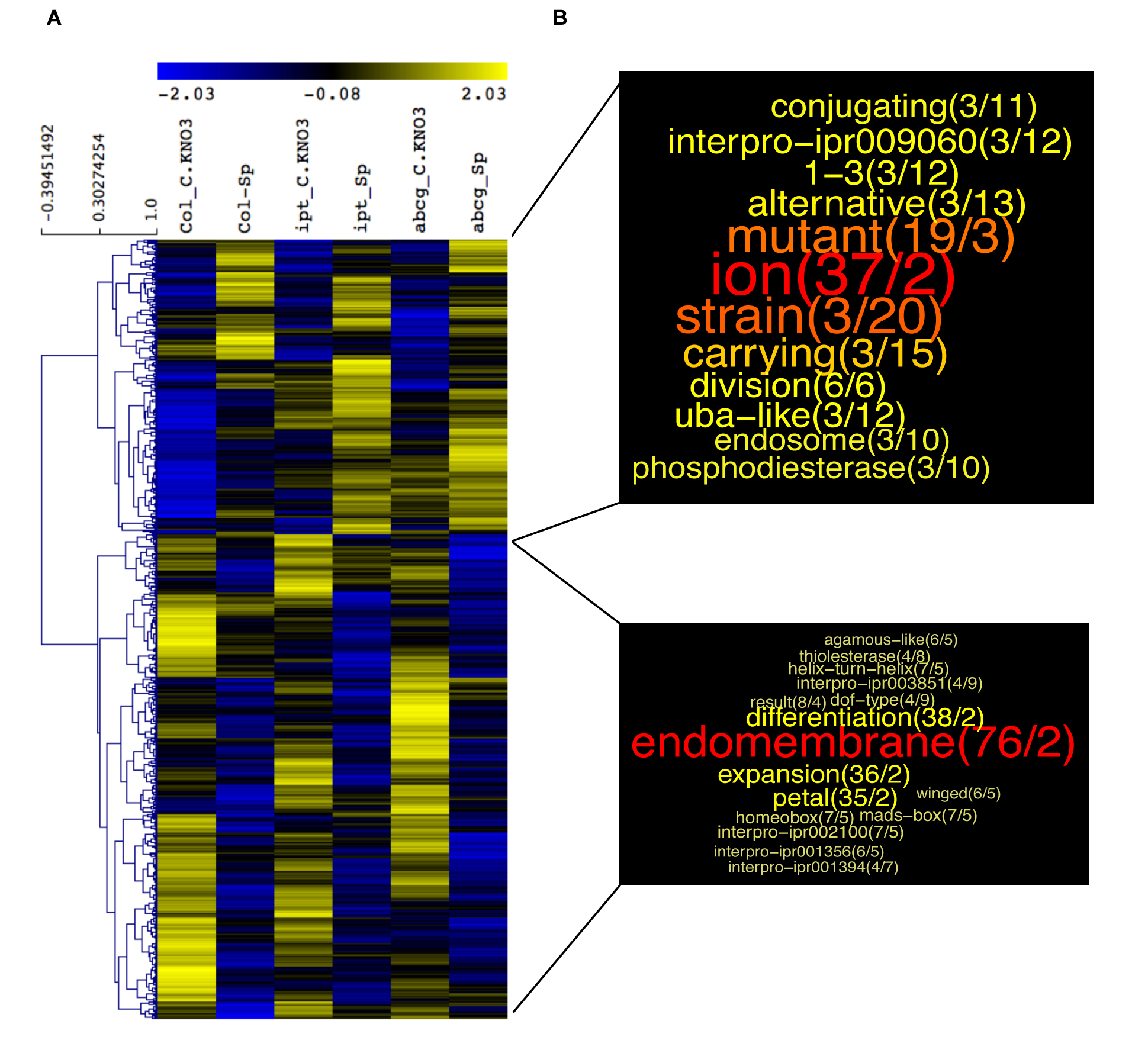
CK-independent shoot transcriptome changes in response to root heterogeneous NO_3_^-^ supply. (**A**) Shoots of Col-0, *ipt3,5,7* and *abcg14* plants transferred for 24 hrs in NO_3_^-^ homogeneous (C.KNO3) or heterogeneous (Split) environment were harvested for transcriptomic analysis. Samples of 4 biological replicates from 4 independent experiments were used to perform the microarray analysis, using Arabidopsis Gene1.1 ST array Strip (Affymetrix GeneAtlas™). Hierarchical clustering of the 669 genes identified as differentially expressed in C.KNO3 versus Split conditions in the 3 genotypes was performed with Multiple Experiment Viewer (MeV) software (http://mev.tm4.org/). (**B**) Semantic enrichment in annotation of genes induced (at the top) or repressed (bottom) in Split condition compared to C.KNO3 in WT shoots, based on a GeneCloud analysis (https://m2sb.org). Beside each term, the first number corresponds to the number of genes containing the term and the second number gives the fold enrichment.

**Supplemental Figure 4.**
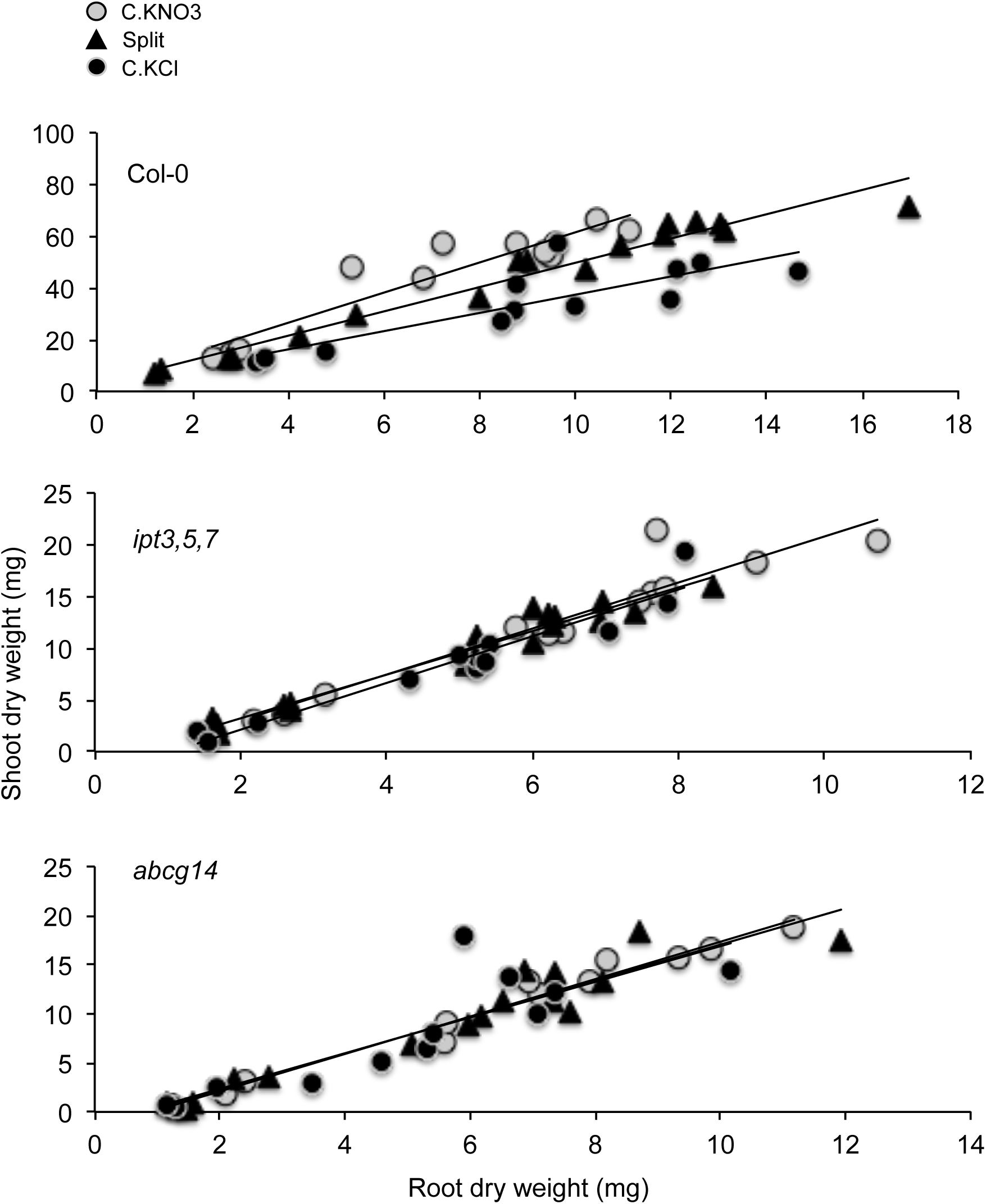
Dynamic of shoot/root ratio in response to N availability is lost in *ipt3,5,7* and *abcg14* plants. Plots display relationship between root and shoot dry biomass from plants treated 4 days in NO_3_^-^ homogeneous (C.KNO3, grey circles), NO_3_^-^ heterogeneous (Split, dark triangles) or N-deprived (C.KCl, dark circles) conditions in the WT, *ipt3,5,7* and *abcg14* mutants. Measurements result from 3 independent experiments. In each experiment, 6 biological replicates by conditions were measured.

**Supplemental Figure 5.**
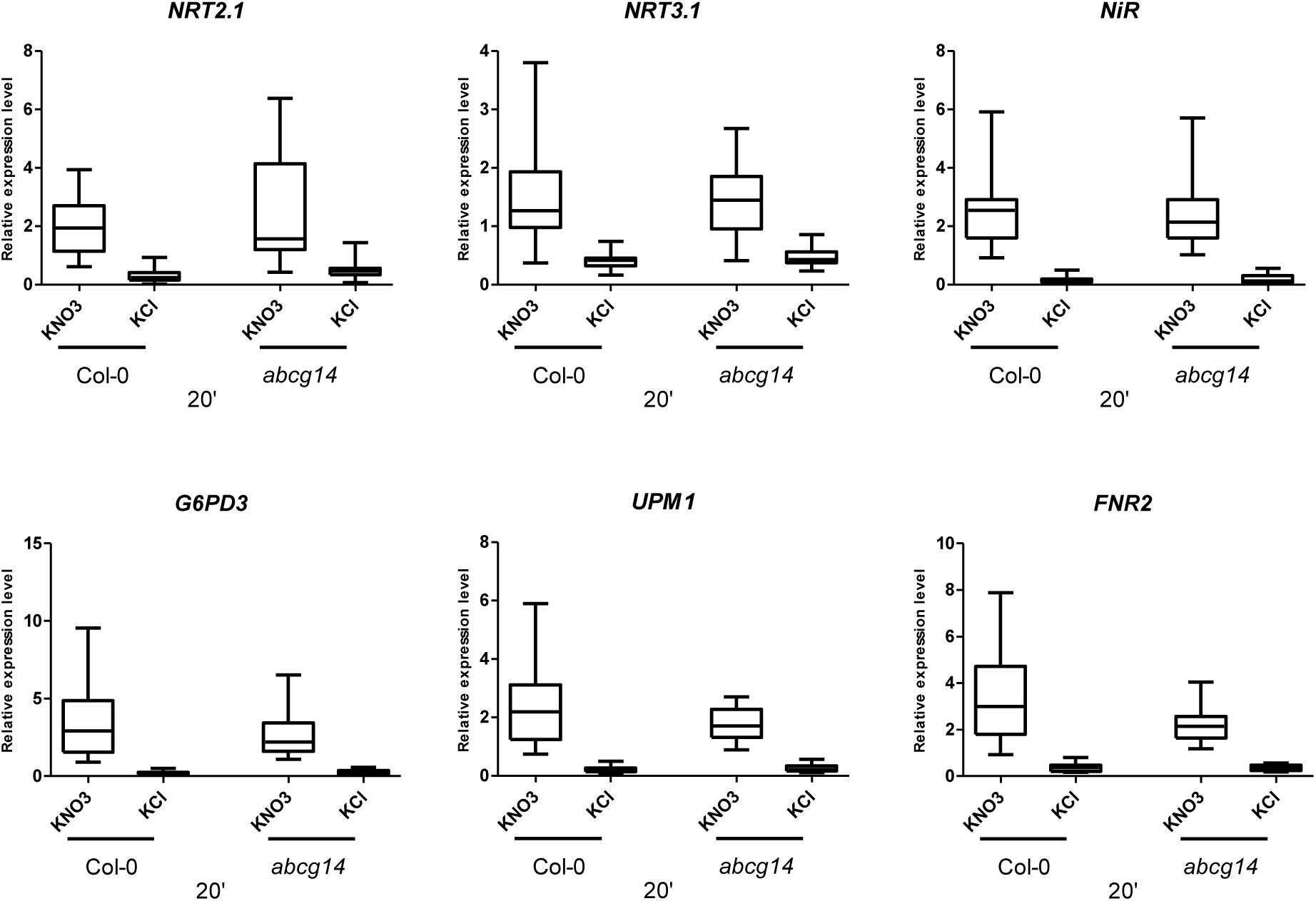
The ability of *abcg14* to trigger the primary NO_3_^-^ response is confirmed in younger plants grown in mini hydroponic system. Boxplots display relative expression level of the 6 marker genes of the primary NO_3_^-^ response: *NRT2.1, NRT3.1, NIR, G6PD3, UPM1* and *FNR2*, in roots of Col-0 and *abcg14*, 20 minutes after transferring the plants in 1 mM KNO_3_ or 1 mM KCl containing medium. Data are means (+/−SE) obtained from 4 independent experiments, including in each 2 or 3 biological replicates that correspond to a pool of 20 to 50 plantlets.

